# DNA Partitioning Modulates Liquid-to-Solid Transitions and the Internal Microstructure of FUS Condensates

**DOI:** 10.1101/2025.06.03.657213

**Authors:** Dea Ilhamsyah, Clement Luong, Seema Qamar, Peter St George-Hyslop, Joel MacKay, Lining Arnold Ju, Tuomas P. J. Knowles, Daniele Vigolo, Yi Shen

## Abstract

Protein liquid-liquid phase separation has recently been recognized as an essential process involved in cellular functions, including transcription, translation, and DNA damage repair. However, a further liquid-to-solid transition (LST) of condensates due to mutation or external stimuli can result in aggregation, sometimes pathological. The unique ability of protein condensates to concentrate and sequester biomolecules is at the heart of the regulatory mechanism, controlling the dynamics and function of the condensates. While the recruitment of essential biomolecules, such as RNA has been studied to have the impact to the condensate formation, how DNA partition can affect the phase behaviour and dynamics of protein condensates is not fully elucidated. In this study, we investigate both the short-term and long-term kinetics of double-stranded DNA partitioning into preformed fresh and aged FUS protein condensates. Confocal imaging shows that DNA partition follows the core-shell diffusion pattern within the condensates. LST slows down and reduces FUS condensates’ ability to recruit DNA but stabilizes the DNA-FUS condensate complex due to the heterogeneous solid network formation. Using the optical technique of Spatial Dynamic Mapping (SDM), we find that DNA partition promotes coalescence and alters the characteristics of the condensates. The partition made the condensates more dynamic in the short term (within minutes) but accelerated LST in long-term incubation (within hours), ultimately leading to an irreversible porous core-shell structure of FUS condensates. Our findings reveal the kinetics of DNA partition during aging and its impact on LST, underlining the modulation of condensate properties by molecule sequestration, shedding light on possible regulation of disease-related LST of biomolecular condensates.

## Introduction

Biomolecular condensates formed by liquid–liquid phase separation (LLPS) provide a dynamic means of spatial organization within cells, enabling the compartmentalization of biochemical processes as membrane-less organelles^1–3^. Condensates exhibit canonical liquid behaviours, including the ability to fuse, drip, deform under shear stress, and rapidly exchange components with their surroundings^1,4,5^. Their liquid-like nature is crucial to support the reversible assembly and disassembly of compartments for biological processes. Biomolecular condensates are inherently multi-component systems, typically comprising of scaffold molecules that drive phase separation and client molecules that are recruited for specific functions. While scaffolds determine the formation and stability of the condensates, clients can be integrated post-condensate-assemblies, sometimes influence the maturation and structure of the condensates. For instance, actin filament structure is determined by the kinetics of actin-binding proteins (ABP) condensates, which over time affect the condensate characteristics^6^.

RNA/DNA-binding proteins (RBP), like TDP-43, hnRNPA1A, α-synuclein and fused-in-sarcoma (FUS) have intrinsic disordered regions and are known to form condensates in cells^7–10^. Their condensates can act as sequesters to accommodate protein-protein, protein-RNA/DNA interactions, involving in the cellular functions, such as stress response, RNA transcription and neuronal functions^10–14^. These proteins are particularly interested due to their association with neurodegenerative diseases^4,10^. Taking FUS protein as an example, its mutations are tightly linked to amyotrophic lateral sclerosis (ALS) and frontotemporal dementia (FTD)^15–17^. These mutations often cause FUS to mislocalize to the cytoplasm, where it undergoes liquid to solid transition (LST) from a dynamic liquid-like state to solid, aggregate-prone forms^10,16^. This pathological aggregation disrupts stress granule function and sequesters essential cellular components, leading to loss of normal nuclear functions, such as DNA repair and RNA processing^18,19^. Together, these dysfunctions contribute to progressive neuronal damage and are characteristic hallmarks of ALS and FTD pathology.

Previous studies have shown that FUS protein can undergo LLPS forming condensates. With aging and physical stimulation, the condensates can undergo further LST forming β-sheet rich solids ^4,5,20^It has been demonstrated that aging of the condensates increases the viscosity, reduce the molecular mobility and the LST is promoted by the condensate surface^21–23^. However, most studies have focused either on single-component systems^20,22,24^ or RNA as a modulator of phase separation^25–27^. Overlooking the complex interplay between scaffold proteins and client molecules. For example, FUS protein forms condensates that recruit DNA fragments^28,29^ and DNA repair proteins such as Ku70/80, NBS1, SFPQ, and 53BP1 (polymerase and ligase)^30^ to facilitate crucial steps in the DNA repair process. However, whether and how the client molecules such as DNA actively contribute to protein condensate physicochemical characteristics is unclear.

In this study, we investigate the spatiotemporal behaviour of FUS condensates with DNA partitioning at different aging stages. We measure the partitioning kinetics, coalescence kinetics, and molecular dynamics by using microscopic technique such as confocal imaging and optical approach such as Spatial Dynamic Mapping (SDM). We found that DNA partitioning into FUS condensates is controlled by the FUS condensates’ characteristics, following a core-shell structure. Furthermore, we demonstrate that DNA recruitment modulates condensate dynamics and internal structures, suggesting that DNA plays an active role in FUS condensate maturation. These findings provide new insights on how to modulate protein condensate phase transitions and prevent irreversible aggregation, shed light on potential treatment and therapies for protein aggregates related diseases.

## Results

### Aging of FUS condensates slows down DNA partitioning

To examine the role of condensates in molecular sequestration, we explored DNA recruitment into FUS condensates. FUS condensates were prepared at a final concentration of 2 μM FUS and 50 mM KCl, following an established protocol^22^. We then aged the FUS condensates for 1 hour (1hr-FUS), 5 hours (5hr-FUS), 24 hours (24hr-FUS), and 48 hours (48hr-FUS) (Fig. 1a). Following condensate formation, we introduced 100 nM of Cy3-tagged DNA at different aging stages (Fig. 1a). Through confocal imaging we observed that DNA partitioned into preformed FUS condensates by propagating from the shell of the condensate to the core (Fig. 1b).

**Figure 1.**
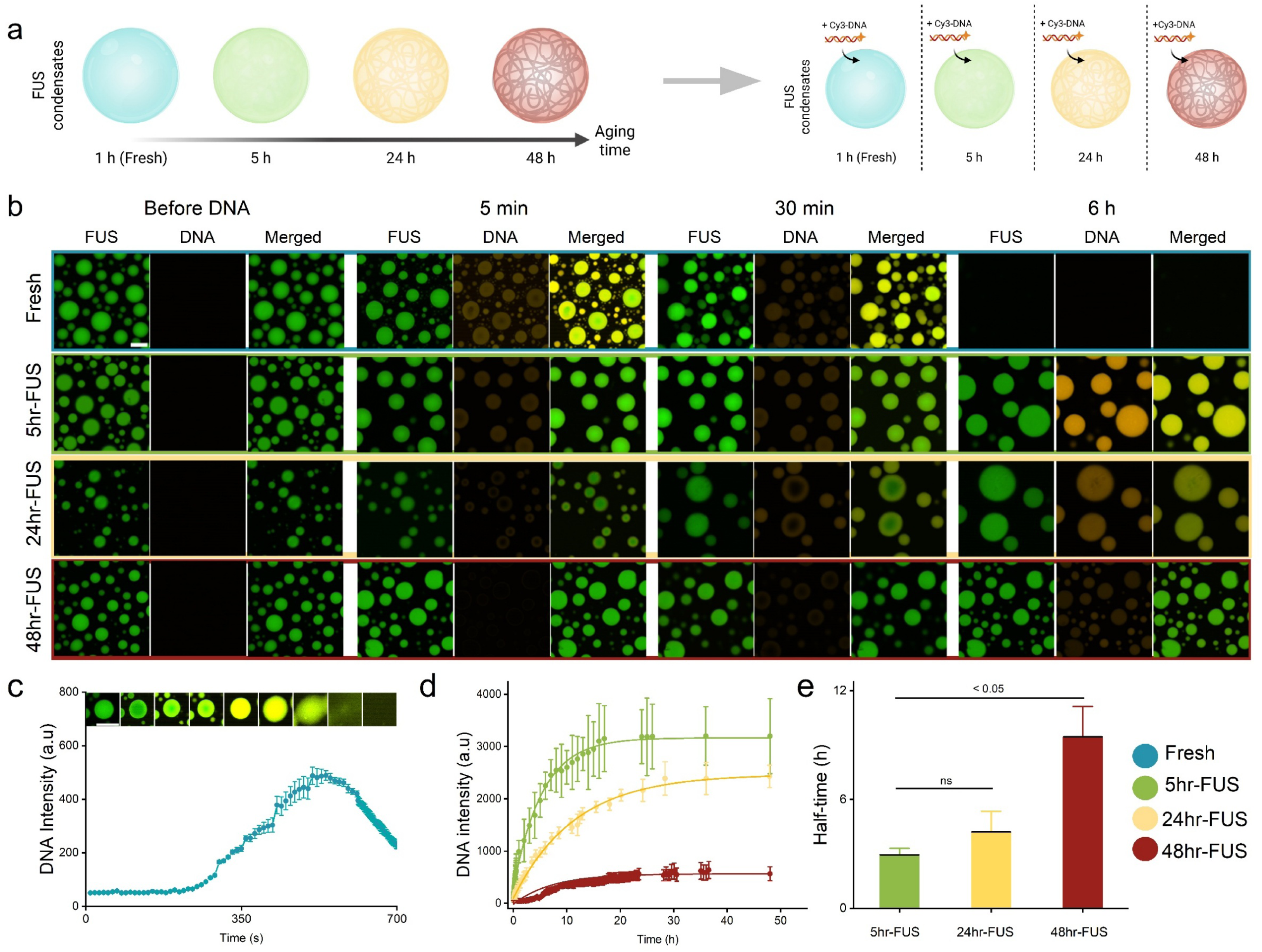
Kinetics of DNA partitioning into different aging stage of FUS condensates. **a**, Schematic diagram of experimental workflow. FUS condensates are aged for different incubation time followed by addition of Cy3-DNA into the solution. **b,** Partitioning of DNA into different age of FUS condensates; Fresh (1hr), 5hr-FUS, 24hr-FUS, 48hr-FUS in different incubation time after introduction of DNA; 5 min, 30 min, 6 hour. (Scale bar = 10µm). c. Raw intensity of DNA partitioning for fresh condensates (n=3) with representative subset images from confocal microscopy observation to show dissolution (Scale bar = 10µm) . **d,** Raw intensity of DNA partitioning into 5hr-FUS, 24hr-FUS, 48hr-FUS (n=3). **e**, Half-time of DNA partitioning (n=3). Half-time is measured based on the time needed to reach half of maximum DNA partitioning intensity inside condensates from different protein batches. T-tests with unequal variance were performed; *P < 0.05; n.s, statistically non-significant values.

To our surprise, freshly formed condensates dissolve upon the partitioning of DNA (Fig. 1b *top row*, c). The fluorescence signal from both FUS and DNA reduce to below detection levels within 700 s, showing that both molecules return to a dispersed state with no further phase separation (Fig. 1c) (Supplementary Video 1). Meanwhile we observed a similar trend of DNA partitioning into aged FUS condensates as seen in freshly formed condensates but without condensate dissolution (Fig. 1b). As DNA is highly charged, we suspect fresh condensates are more sensitive to the electrostatic changes due to the high internal molecular dynamics, while semi-aged (5-24 hours) condensates are still flexible to recruit DNA molecules but more stable to resist the dissolution due to the fluctuation of the charge (Supplementary Video 2).

Interestingly, after 30-minutes of observation, the DNA partitioning for 24hr-FUS is mostly at the shell of the condensates unlike for 5hr-FUS, for which the DNA is already at the core, indicating there’s a difference of partitioning kinetics due to aging of FUS condensates. To confirm this, we continued monitoring DNA partitioning until the DNA fluorescence signal reached a plateau. Average intensity of DNA signal is measured from at least 3 condensates as a function of time (Fig. 1d). Results have shown that DNA signal increases at all aging states and plateaus at 17-25 hours depending on the aging. To be noted, there is no decrease in FUS fluorescence intensity with the partitioning of DNA, indicating that there is only influx of DNA into FUS condensates. During the recruitment, 5hr-FUS condensates have the highest DNA intensity, followed by 24hr-FUS and then 48hr-FUS (Fig. 1d). Previous study has shown that FUS condensates can undergo LST within 48 hours^22^. Our findings suggest this aging induced LST hinders the ability of FUS condensates to take up DNA. The reduced ability may result from the loss of available heterotypic (FUS-DNA) interacting sites, as charged and aromatic residues becoming locked into stable hydrogen bonded β-sheets^4,31,32^.

Furthermore, we noticed there were differences in the rate of DNA partitioning across aging stages. To quantify these differences, we measured the half-time of DNA partitioning as the time needed for DNA to reach half of its maximum concentration in the condensates. Condensates formed from different experiments were used to assess replication. The results indicated that 48-hour-aged condensates recruit DNA approximately two times slower than 5-hour and 24-hour samples (Fig. 1e). These findings support the hypothesis that LST progressively impairs the ability of condensates to recruit DNA. We propose that in the more solid-like 48-hour-aged FUS condensates, the reduced partitioning efficiency is due to decreased molecular mobility, and increased internal viscosity and elastic modulus of FUS ^24^, which together hinder both the diffusion of DNA and the FUS–DNA interactions.

### Diffusion pattern of DNA partitioning into FUS condensates during aging

Our data has shown a distinct diffusion pattern of DNA within FUS condensates. To gain insights into the effects of LST-induced structural heterogeneity, we measured the DNA diffusion (estimated based on its fluorescent intensity) across five distinct regions within the condensates over time, from the shell to the core (Fig. 2a). It is important to note that we measured the recruitment kinetics of condensates with similar sizes (8-10 µm in diameter) to avoid bias in volume differences, while still capturing the variation in diffusion characteristics between the shell and core regions.

**Figure 2.**
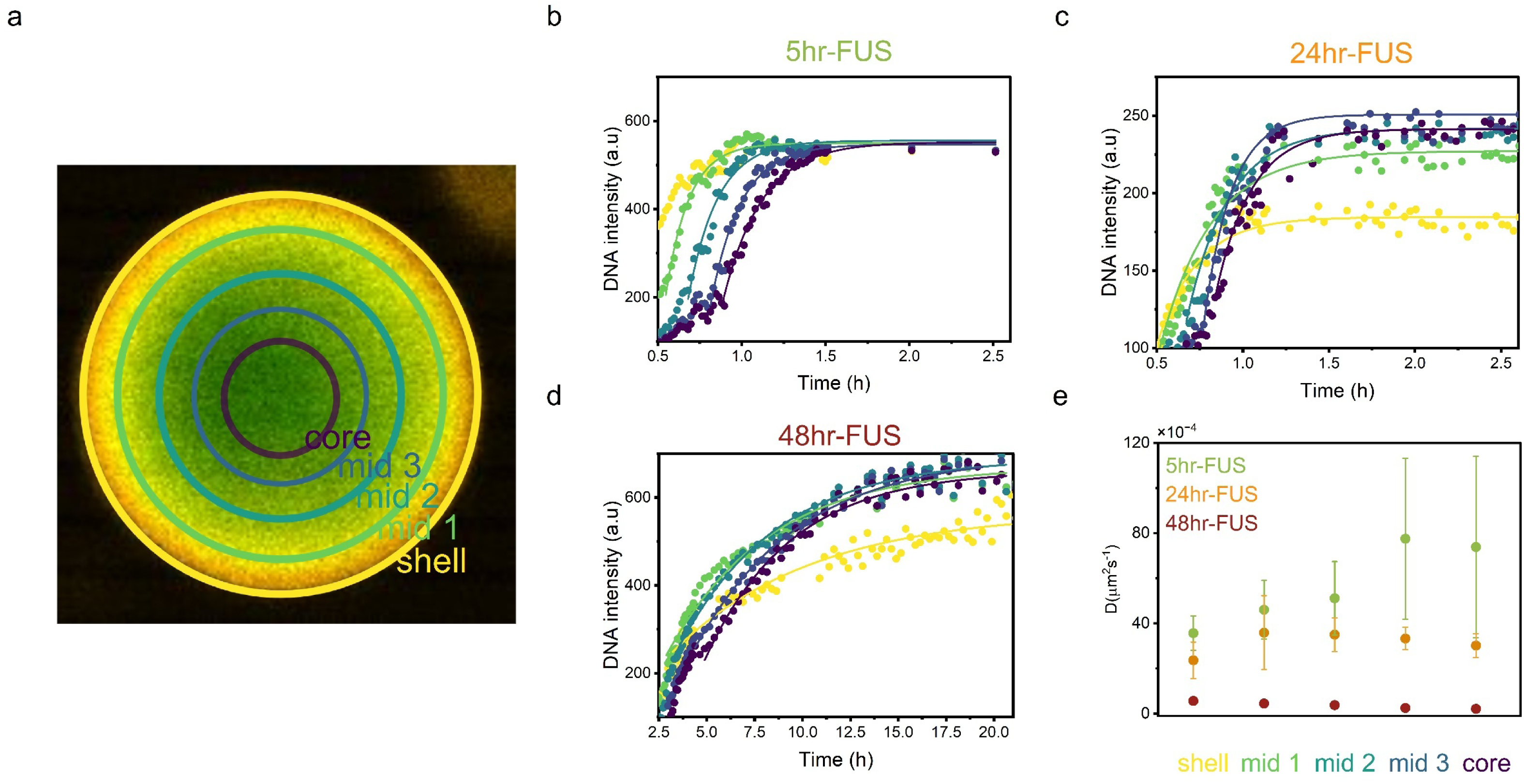
Diffusion pattern of DNA inside aging FUS condensates. **a,** Merged confocal figure for visualization. FUS condensates are divided into 5 regions of interest: Shell, Mid 1, Mid 2, Mid 3 and Core. DNA intensity data were obtained from average intensity of DNA fluorescence signal for the ROI. **b,** Core-shell DNA intensity plot in 5hr-FUS.**, c**, Core-shell DNA intensity plot in 24hr-FUS. **d,** Core-shell DNA intensity plot in 48hr-FUS. **e,** Diffusion coefficient (D) of DNA inside FUS condensates measurement in different ROI at three aging stages (5hr-FUS: green, 24hr-FUS: orange, 48hr-FUS: red). n=3. DNA diffusion coefficient in FUS condensates is obtained from three independent repeated experiments.

Our results show that in semi-aged 5-hour and 24-hour FUS condensates, DNA reaches the core of the condensate within 2.5 hours (Fig. 2b, c), while 48-hour FUS condensates require more than 15 hours (Fig. 2d). These results are consistent with previous data, indicate that the diffusive behaviour of DNA varies across different aging stages. Interestingly, for 24-hour and 48-hour condensates the plateau concentration of DNA at core is higher than at the shell, showing a core-shell pattern. We suspect this is due to the greater recruitment capacity of the condensate’s dynamic core.

We further calculated the diffusion coefficient of DNA in the five regions inside FUS condensates (Fig. 2e). The diffusivity of DNA in 5-hour FUS condensates ranged from 35 µm²/s at the shell, with an increase to 74 × 10⁻⁴ µm²/s in the core area. Similarly, in 24-hour FUS condensates, DNA diffusivity range of 23 to 30 × 10⁻⁴ µm²/s, showing only a subtle difference of diffusion from shell to core areas. Notably, in 48-hour condensates, this core-shell diffusion pattern was no longer observed, and the overall diffusivity dropped significantly to 5 × 10⁻⁴ µm²/s, an order of magnitude slower than in the earlier aging stages. This suggests that aged condensates slow down the diffusion of client molecules due to the increase of the viscosity^24,33^.

Despite previous studies having shown that there is a distinct characteristic within the core-shell area of condensates^20–22^, we suspect that the absence of a core-shell DNA diffusive gradient is because of the short hydrodynamic radius of the DNA. Our model DNA is relatively small with 207 base pairs; therefore, the formation of a gradient gel-like network from shell to core was not sufficiently dense enough to obstruct DNA movement and affect its diffusion behaviour.

### DNA partitioning leads to SLT of FUS condensates

Building on our observations of altered DNA diffusion dynamics, we next examined how DNA partitioning changes over time and whether it influences the physical properties of FUS condensates. To investigate how condensate stability responds to increasing DNA levels at different aging stages, we tested a range of DNA concentrations (50–300 nM), based on previous findings that RNA influences condensate dynamics in a highly concentration-dependent manner^27^.

Confocal imaging was performed after 24 hours of incubation of the mixture of the condensates and DNA, allowing the system to reach near-equilibrium (Fig. 3a). We found the addition of high concentrations of DNA (100-300 nM) dissolved freshly formed FUS condensates (Fig. 3a). In contrast, aged condensates exhibited greater resistance to high DNA concentration and underwent a solid-to-liquid transition (SLT), leading to coalescing events. Interestingly, 5hr-FUS only dissolve at DNA concentration higher than 200 nM. Meanwhile, 1wk-FUS condensates remained even with high DNA concentration. Some of the 1-week-aged condensates, particularly the small ones, retained the ability to recruit DNA (Fig. 3a, red arrows). This might be the result of the SLT of FUS condensates at the highest concentration of DNA. Together, these results suggest that FUS condensates remain in a liquid-like state up to 48 hours of aging, beyond which they acquire solid-like characteristics that mostly resist both dissolution and partition of DNA.

**Figure 3.**
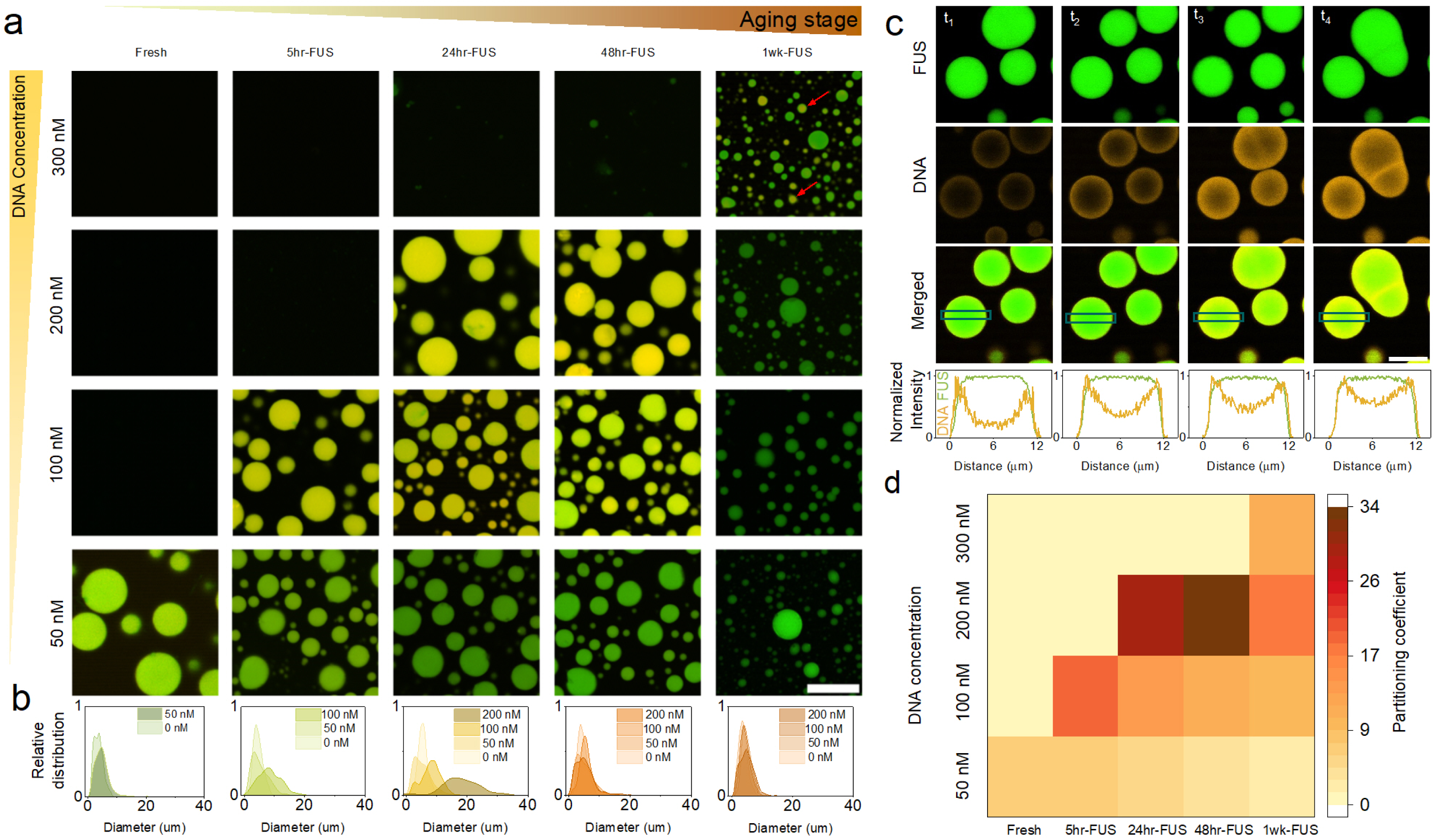
Effect of DNA partitioning on the phase diagram of aging FUS condensates. **a,** Merged fluorescence images of GFP-FUS and Cy3-DNA channels. Representative images are showing DNA (50-300 nM) partitioning in FUS condensates at different aging stages (Fresh, 5hr-FUS, 24hr-FUS, 48hr-FUS and 1wk-FUS). Red arrows show partial partitioning of DNA in 1wk-FUS. Scale bar = 20 µm. **b,** FUS condensates size distribution at different DNA concentrations. **c,** *Top*. Confocal fluorescence images of the coalescence of FUS condensate upon DNA partitioning (Green: GFP-FUS, Orange: Cy3-DNA, Green + Orange: merged). Scale bar = 10 µm. *Bottom*. Plot intensity from GFP-FUS and Cy3-DNA channels. **d,** Heatmap of DNA partition coefficient (K_p_) in different concentration of DNA (50-300 nM) and different FUS aging stages. The images are representative of the observed reproducible behaviour from at least three test replicates for each corresponding DNA concentration and FUS aging time.

We next analysed condensate size distributions across varying DNA concentrations and aging stages (Fig. 3b). As expected, increasing DNA concentration leads to larger condensates in fresher samples due to coalescence, while size shifts are minimal in aged condensates, again, likely due to the formation of more solid-like networks in 1-week-old FUS. Notably, coalescence occurred once DNA began to be recruited into condensates (Fig. 3c). Coalescence times increased by two orders of magnitude for 5- and 24-hour FUS, and by three orders for 48-hour FUS (Supplementary 1a). Condensate fusion followed an exponential relaxation trend in aspect ratio, from which we calculated relaxation times of 55 s for 5-hour, 73 s for 24-hour, and 1986 s for 48-hour FUS (Supplementary 1b). Where no coalescence was observed in single-component FUS condensates beyond 48 hours of aging. These findings are consistent with previous studies, where single-component fresh FUS condensates had relaxation times ranging from milliseconds to 10 seconds, while aged condensates (<30 min) showed relaxation times of approximately 30 seconds^24^.

Previous study suggests that the coalescence of condensates can rearrange the location of DNA at the condensate interface^34^. The primary driving force behind condensate coalescence is the reduction of interfacial free energy^4,35^, with droplets fusing to form larger ones due to the energetically favourable decrease in surface area. This is aligned with our observation that the coalescence of FUS condensates starts before DNA reaches the core (Fig. 3c, Supplementary 1), suggesting that molecular influx immediately drives coalescence by altering condensate composition, surface tension, and internal dynamics.

Finally, to assess the impact of the aging process, we measured the partition coefficient of DNA at different concentration and aging stages. Briefly, the partition coefficient (Kp) is the ratio of the client molecule concentration in the condensed phase to the dilute phase^36,37^. Results show that, for the low DNA concentrations, the partition coefficient decreases as condensates age (Fig. 4d). However, for the high DNA concentrations, aged condensates recruit and retain the DNA molecules, while fresher condensates dissolve into the disperse state, in agreement with the previous results.

**Figure 4.**
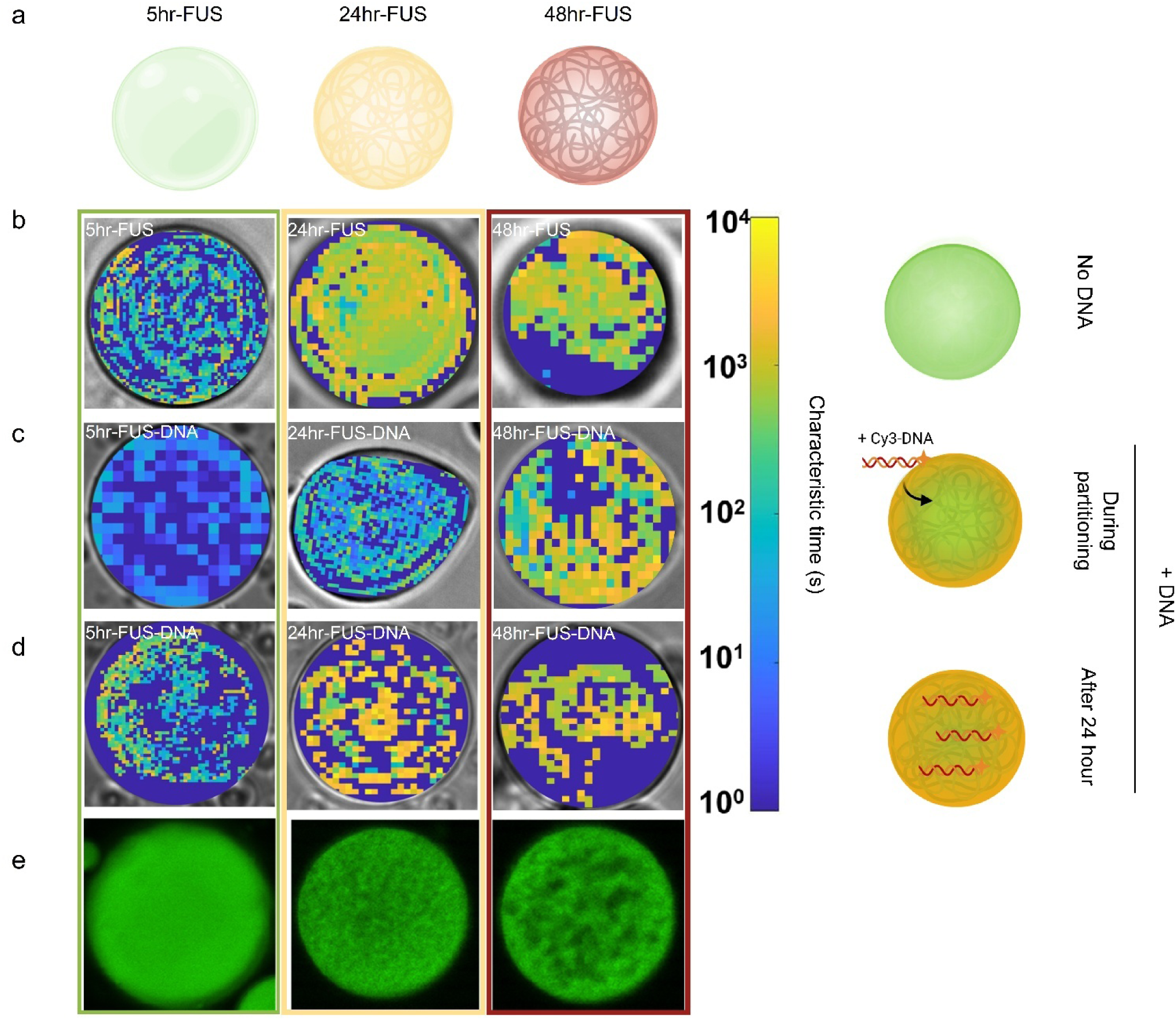
SDM analysis of FUS and FUS-DNA condensates in different stage of aging. **a,** Illustrations of FUS condensates at different aging stages. 5hr-FUS (green), 24hr-FUS (yellow), and 48hr-FUS (red) **b**, SDM analyses before DNA partitioning. **c**, SDM analyses during partitioning, after DNA reaches the core of condensates. **d**, SDM analyses of FUS-DNA condensates after 24-hour incubation, when DNA partitioning reaches the saturation inside condensates. Side illustration represents the DNA introduction to FUS condensates at different time points. **e.** Representative confocal images of porous FUS condensates (GFP signal alone).

These findings suggest that while aging reduces the ability of condensates to actively recruit client molecules from the surrounding environment, aged condensates develop a stable microenvironment that retains and concentrates client molecules. In disease-related contexts, a lower ability of FUS to recruit client molecules might result in compromised cellular functions. On the contrary, condensate aging may also play a role in preserving key biomolecular interactions or buffering undesired components in high molecular flux conditions (e.g., stress, overexpression, high molecular signalling exchange)^38–41^.

### DNA partitioning increases the dynamics of FUS condensates in the short term but accelerates LST in the long term

Moving forward, we focus on the effects of DNA partitioning over different time scales on the dynamics within FUS condensates. Specifically, we aim to investigate the effects of short-term DNA partitioning and the long-term outcomes once DNA partitioning plateaus. Here, we probed the dynamics within the condensates using a bespoke previously reported Spatial Dynamic Mapping (SDM) optical technique to reveal the material properties within the condensates at high spatial resolution. Briefly, this method enabled us to measure the dynamics of molecules at specific locations within a condensate by analysing the spatial variance in real space over time^22^. A short characteristic time indicates liquid-like behaviour, while a long characteristic time corresponds to solid-like behaviour.

We investigate the changes in dynamics within 5hr-, 24hr- and 48hr-FUS condensates in the presence of 100 nM DNA (Fig. 4a). Using SDM, we find that FUS condensates alone become more solid-like during aging, consistent with previous studies^20,22^(Fig. 4a). Subsequently, we performed SDM analysis upon DNA partitioning at different aging stages of the condensates. The analysis is performed after the DNA reaches a uniform intensity from shell to core (2.5 – 6 hours depends on the aging stage). The results reveal that all condensates become more dynamic after the recruitment of DNA (Fig. 4c). This finding aligns with other studies, which have shown that the exchange of molecules enhances the overall dynamics of condensates through molecular rearrangement within the dense phase^42^.

Next, we examined the dynamics of the condensates after 24 hours aging of FUS-DNA condensates. All 5hr-FUS-DNA, 24hr-FUS-DNA and 48hr-FUS-DNA condensates became more solid-like. Interestingly, we observe a greater degree of heterogeneity in dynamics (Fig. 4d) resembling a porous network, that is also demonstrated by the confocal images of the condensates (GFP signal alone) (Fig. 4e). From our results, it has shown that although DNA partitioning can dissolve and increase the dynamicity of the condensates, the LSTs still occur during aging after the recruitment of molecules. Meanwhile, the partitioning of DNA might alter the irreversibility of LST of condensates, resulting in more uneven network formation within the condensates.

### DNA partitioning introduces irreversible microstructural changes of FUS condensate

To probe internal spatial reorganization effects from DNA partitioning, we performed a dissolution assay. Briefly, we added 1 M KCl to FUS-DNA condensates after a 24-hour incubation time (Fig 5a). This incubation time is chosen to ensure DNA partitioning reaches a near plateau.

**Figure 5.**
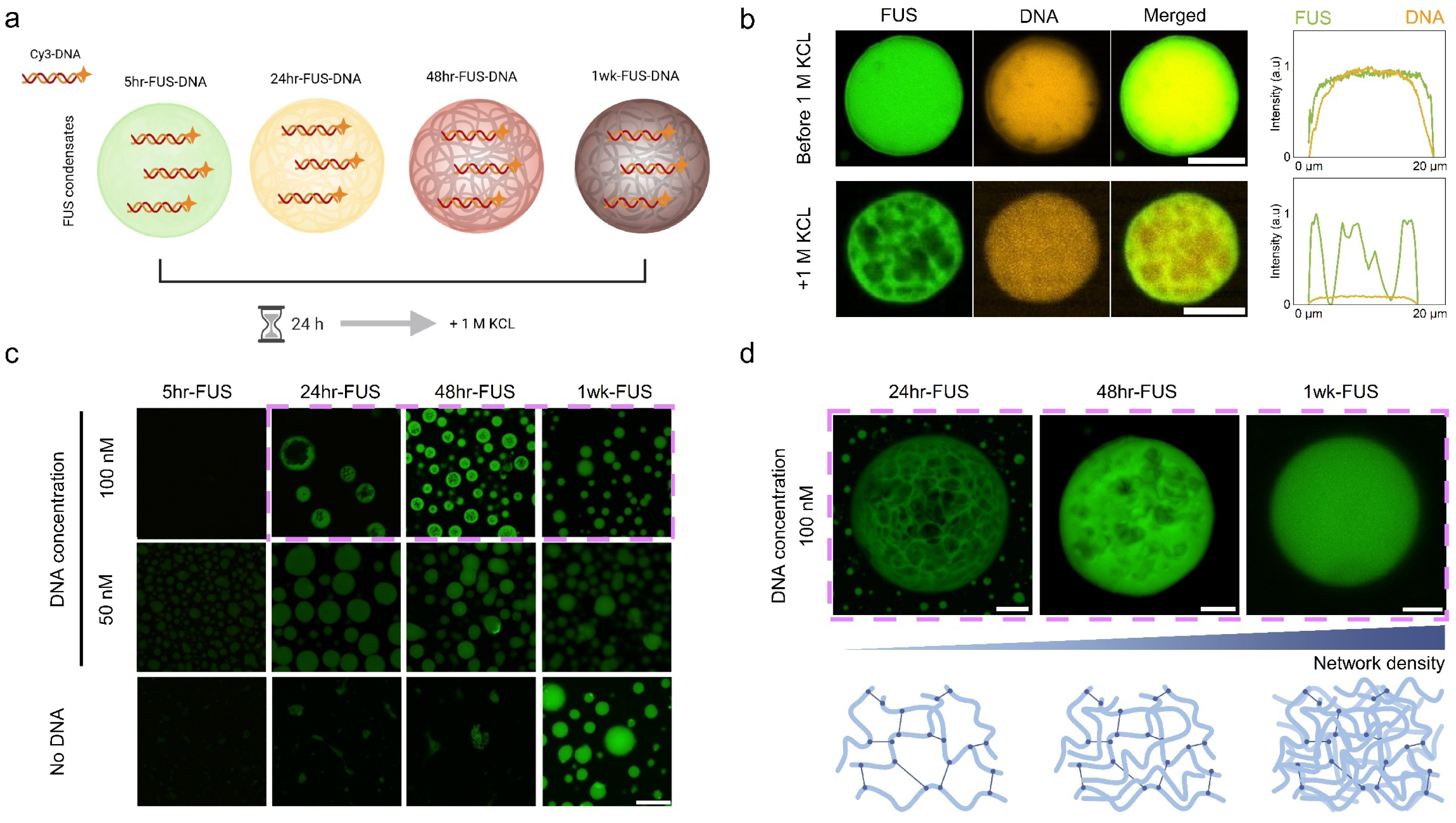
Dissolution experiment of FUS-DNA condensates. **a**, Schematic diagram of aging and dissolution of condensates **b**, Representative fluorescence images and intensity profiles of 24hr-FUS-DNA condensate. *Top row*, before dissolution. Images are showing DNA accumulate at the core of condensates, *Bottom row,* after dissolution. Porous FUS protein remains while DNA signal is low, showing most DNA released back to the dispersed state (Scale bar = 5 µm). **c**, Overlay fluorescence images of DNA-FUS condensates after dissolution. *Top row*, 100 nM DNA is added. *Mid row*, 50 nM DNA is added. *Bottom row, control* group of FUS condensates aged without DNA (Scale bar = 10 µm) **d**, *Top* 3D projection of Z-stack confocal images of condensate internal structure after dissolution (100 nM DNA partitioning, Scale bar = 5 µm). *Bottom* Illustration of network density of condensates at different aging stages.

As aging progressed, we observed that DNA no longer distributed evenly throughout the condensates but instead localized preferentially in the core of 24hr-FUS-DNA condensate (Fig 5b). The rearrangement of DNA within the condensates, likely results from the development of a core-shell architecture during aging^21–23,43^. As the shell presents a more crosslinked and solid-like structure, it loses the ability to recruit DNA molecules. In contrast, the more dynamic and interactive core promotes DNA recruitment and retention. Following this, a dissolution assay was performed (Fig. 5b).

Upon the addition of KCl solution, the condensates dissolve quickly with most of the fluorescent signal dispersed into the diluted phase (Supplementary Video 3). A porous and heterogeneous FUS network is left with a very small amount of DNA evenly distributed through the condensate (Fig. 5b bottom row). This experiment suggests the irreversible DNA-protein covalent binding is minimum in the protein-DNA condensate and does not take part in the solid network formation. Instead, there are transient bindings between DNA and protein molecules, influencing the condensate phase behaviour.

Furthermore, we compared the dissolution results without and with the presence of different concentration of DNA (0, 50, 100 nM) (Fig. 5c). Interestingly, with 50 nM DNA partitioned, FUS condensates barely dissolved at all aging stages (Fig. 5c mid row). Meanwhile, with 100 nM of DNA partitioned, 5hr-FUS condensates dissolved but the semi-aged condensates (24hr- and 48hr-FUS) showed porous solid network structure and a solid shell after dissolution (Fig. 5c top row). Comparatively, with the absence of DNA partitioning, all FUS condensates dissolved except for 1wk-FUS condensates (Fig. 5c bottom row). This result indicates that without DNA partitioning, FUS condensates remained in high-viscosity liquid as opposed to an irreversible porous solid network. In the presence of DNA, transient interactions occur between DNA and protein. We suspect, at a high concentration, DNA saturate the active binding sites and disrupt the protein-protein interactions, resulting in formation of solid structures. In contrast, at a lower concentration, DNA molecules available were not enough to occupy all the active binding sites of FUS. It possibly screens the charge and enhance the protein-protein interactions, accelerating condensate LST.

Finally, using 3-D projection confocal images, we demonstrate zoomed-in porous solid network within the condensates after dissolution (Fig. 5d). With the DNA partitioning at different aging time, the FUS network density changes. 24-hr FUS shows an irreversible porous core-shell structure after dissolution (Supplementary Video 4), while 48-hr FUS exhibits visibly smaller pore sizes (Supplementary Video 5). Finally, 1 wk-FUS shows a dense shell with no visible pores (Supplementary Video 6). These changes in porosity and morphology suggest that recruitment of nucleic acid can modulate the internal microstructures of FUS condensates.

## Discussions

In this study, we investigated the dynamic interplay between DNA and FUS condensates during aging, revealing how client molecules can modulate the condensate phase behaviour and structural organization. Upon the addition of DNA, over a short timescale, fresh condensates rapidly dissolve, while aged condensates retain their structure and recruit DNA over time. The partitioning of DNA propagates from shell to core, with DNA molecules accumulated at the core of the condensates at last.

Using diffusion kinetics analyses, we found that the degree of DNA partition decreases with condensate’s age. A core-shell diffusion pattern emerges at semi-aging stages. This pattern fades in aged condensates as a solid gel formed across the whole condensate. Our findings further demonstrate that DNA partition has dual effects on FUS condensates: while short-term DNA recruitment can enhance dynamics and promote coalescence via solid-to-liquid transitions (SLT), in the long term, the partition accelerates condensate solidification. Interestingly, DNA appears to accumulate more in the core than the shell in aged condensates, likely due to increased viscosity and solid network in the shell that limit the partition. The formation of an irreversible porous network of FUS emerged while most of the DNA was released into a dispersed phase. It has demonstrated that, depends on the concentration, client molecules can drive internal reorganization and alter the microstructure of condensates.

Together, these findings suggest that DNA plays a dual role: it transiently enhances condensate fluidity at early stages but ultimately contributes to the maturation and solidification of FUS condensate over time. This highlights that DNA is not a passive client but act as a modulator of condensate phase behaviour, capable of shaping both phase stability and material properties depending on two factors: client molecule concentration and aging time. This work expands our understanding in how nucleic acid-protein interactions may contribute to either physiological or pathological liquid-to-solid transitions of biomolecular condensates. Our results provide a range of insights into the fundamental principles governing condensate phase behaviour and offer a foundation for developing condensate-based therapeutic strategies.

## Methods

### DNA preparation

Labelled nucleosomal DNA was prepared by large-scale PCR (5 mL) amplification of the 0w60 DNA sequence with Cy3-labelled forward primer and reverse primer supplied by Integrated DNA Technologies. The PCR was carried out using Phire Hot Start II DNA Polymerase under standard conditions (98°C for 20 s, 54°C for 20s, 72°C for 10 s, 35x cycles).

The PCR product was precipitated by the addition of sodium acetate (300 mM), followed by 3 volumes of ethanol. The sample was allowed to precipitate (20 min on ice), then centrifuged at 15000 g, 4°C for 30 min. The supernatant was removed and the pellet dissolved in 1x TE buffer (10 mM Tris-HCl pH 7.5, 1 mM EDTA), and mixed with an equal volume of phenol:chloroform:isoamyl alcohol (25:24:1). The aqueous layer was taken and washed with an equal volume of chloroform, then precipitated with the addition of ammonium acetate (2.5 M), followed by 1 volume of isopropanol. The pellet was washed with 70% ethanol and then dissolved in 1x TE buffer and 40% (w/v) sucrose The sample was then loaded onto a 5% polyacrylamide TBE gel for native gel electrophoresis. The band containing 0Cy3w60 DNA was excised and subjected to electro-elution at 150 V for 25 min, then precipitated by the addition of sodium acetate (300 mM), followed by 3 volumes of ethanol. The DNA was redissolved in 1x TE buffer, and the concentration of DNA was measured using a Nanodrop® 1000 spectrophotometer. DNA sequence is available on supplementary data (See Supplementary 2).

### Protein Expression

FUS protein was produced using baculovirus expression system vector, pACEBac2, mentioned in previous work^5^. Briefly, the purified recombinant bacmid encoding FUS was transfected into Sf9 insect cells, where protein expression occurred for 4 days following infection with the baculovirus containing the recombinant bacmid. Afterward, the cell culture was harvested by centrifugation at 4,000 r.p.m. for 30 minutes. The resulting cell pellet was then homogenized in lysis buffer composed of 50 mM Tris, 1 M KCl, 0.1% CHAPS, 1 mM DTT, and 5% glycerol, adjusted to pH 7.4. Following lysis, the sample was centrifuged at 40,000 r.p.m. in an ultracentrifuge to eliminate cell debris. The supernatant was collected and subjected to a three-step purification protocol, which included a nickel nitrilotriacetic acid affinity column, an amylose affinity column, and a final size exclusion chromatography polishing step using a buffer containing 50 mM Tris, 1 M KCl, 1 mM DTT, and 5% glycerol at pH 7.4. The final protein purity achieved was greater than 95%.

### Sample Preparation for DNA Partitioning

All experiments were conducted in PDMS (Polydimethylsiloxane) wells attached to a glass coverslip. The wells were created using a PDMS slab (±1 cm thick) and were punched with a 1.5 mm biopsy punch (KAI Biopsy Punch 1.5 mm). This well size was selected to minimize sample volume, reduce the impact of slow diffusion associated with larger volumes, and slow evaporation rate compared to shallower wells.

To prevent wetting of the condensates, the coverslips were cleaned and pretreated with plasma treatment and PEG-Silane (3-Methoxy(polyethyleneoxy) 9-12[propyltrimethoxysilane]) (Gelest, SIM6492.72). First, the coverslips were soaked in acetone for 1 hour, rinsed with MilliQ® water, and dried with nitrogen gas. The PDMS wells were then plasma-bonded to the coverslip using a plasma cleaner (Zepto, Diener Electronic Plasma Cleaner). Afterward, a 1:1 (v/v) mixture of PEG-Silane and ethanol was pipetted to cover the bottom of the wells, which were then heated on a 65°C hot plate for 15 minutes. Any remaining solution was washed away with MilliQ® water, and the wells were dried using nitrogen gas and left to dry overnight.

### Confocal Microscopy

All confocal images were taken using Nikon Ti2 Eclipse Confocal Microscope equipped with 20x objective lens magnification with 10x zoom. 512 x 512 pixels images were taken over time for 24 hours, with exact laser power: 0.1 for Laser 488 nm (499 nm - 541 nm emission range GFP) and 4.0 for Laser 561 nm (571 nm - 625 nm emissions range Cy3). Z-scans of images were projected using NIS-element AR.

### Partitioning Dynamics and Diffusivity Measurement

The diffusivity of DNA within protein condensates was measured from time-series confocal images. Five condensates, each with an average size of 8-10 µm in diameter, were selected for analysis. To minimize measurement errors due to condensate drift, drift correction was applied to the confocal images. Specifically, the FUS-GFP channel was used as a reference to determine the drift parameters, which were then applied to correct the DNA signal. The drift correction code is available in NanoPyx^44^. Following drift correction, image analysis was performed using FIJI^45^. For the analysis of recruitment at different positions within the condensates, each condensate was divided into five concentric regions of varying radius (see Fig. 2a). From each area, *j*, and from each time point, we extracted the radial average of the fluorescence signal intensity, 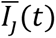. By plotting 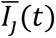 for each area we were then able to extract the characteristic diffusion time, *τ*_*D*_, knowing that mass diffusion follows an exponential dynamic

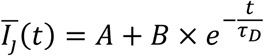

and then we extracted the diffusion coefficient, *D*, based on the equation

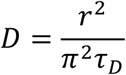

Where *r* is the radius of the condensate.

It is important to note that, as DNA diffuses from outside the condensate towards its centre, the exponential behaviour is delayed going from the shell to the core as shown in Fig. 2b-d.

### Coalescence Dynamic

Images were taken using confocal microscope with 20x objective lens magnification with 10x zoom. Coalescence dynamics were measured by the characteristic relaxation time, as previously described in the referenced study^46^. The aspect ratio of condensate was calculated by measuring the fitting an ellipse to the shape and calculating *a*⁄*b*, where *a* and *b* are the long and short axes of the ellipse.

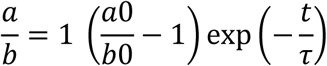

where *t* is time, *τ* is the characteristic relaxation time, and *a*0⁄*b*0 is the initial aspect ratio before coalescence.

### Partition Coefficients

Partition coefficients (K_p)_ were calculated from fluorescence confocal microscopy images with the exact same laser power for all variations of aging stages (Laser 488 nm, Power: 0.1 for GFP and Laser 561 nm, Power: 3.0 for Cy3). K_p_ were defined as the raw average intensity of DNA inside FUS condensate compared to the average intensity of background value.

### Data analysis

Condensate diameters were quantified from confocal images using a custom macro script developed in ImageJ. The partitioning half-time value were calculated from average intensity of DNA within each condensates overtime. The half-time were defined as time needed to reach half of the maximum fluorescence intensity based on other study^27^. Graphs and statistical analyses were done using Origin 2024b.

## Author contributions

Y.S. conceived the study. D.I. and Y.S. designed and conducted the experiments with advice from D.V. D.I., Y.S. and D.V. analysed and interpreted the data. J.M, C.L, P.S.G., S.Q, L.A.J., T.P.J.K., D.V. and Y.S. co-wrote the paper with input from all the authors.

## Acknowledgments

We acknowledge financial support from the European Research Council through the ERC Grant Di-ProPhys to Prof. Tuomas Knowles (Agreement 101001615). We acknowledge the support from Australian Research Council Discovery Early Career Researcher Award (DE230100837) for Dr. Yi Shen.

---

**Supplementary 1.**
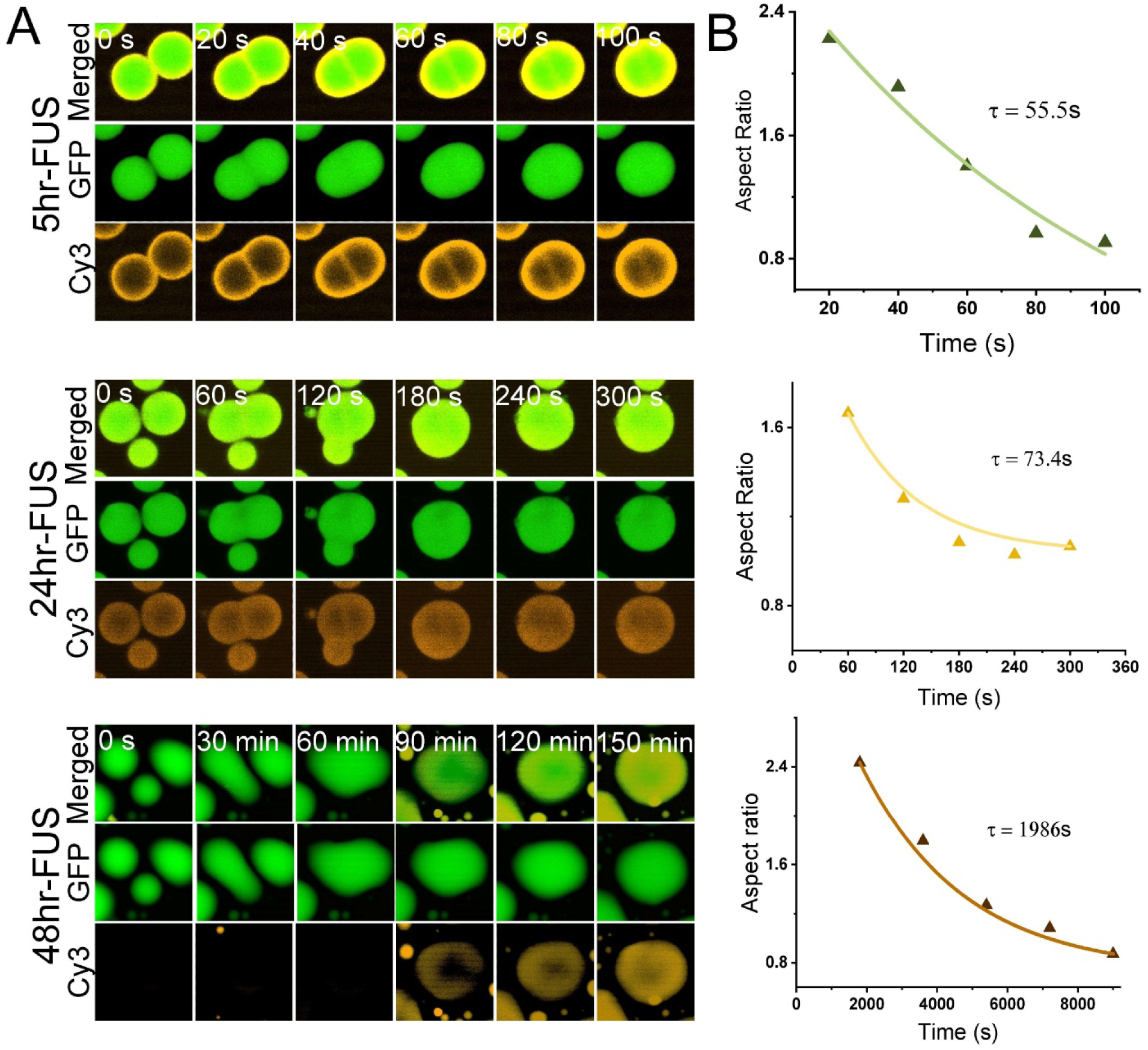
Coalescence dynamic of DNA partitioning into aging FUS condensates A Fluorescence microscopy images from coalescence dynamic across aging FUS condensates in different time scale. B Relaxation time measurement; 5hr-FUS (top), 48hr-FUS (middle), and 48hr-FUS (bottom).

**Supplementary 2.**
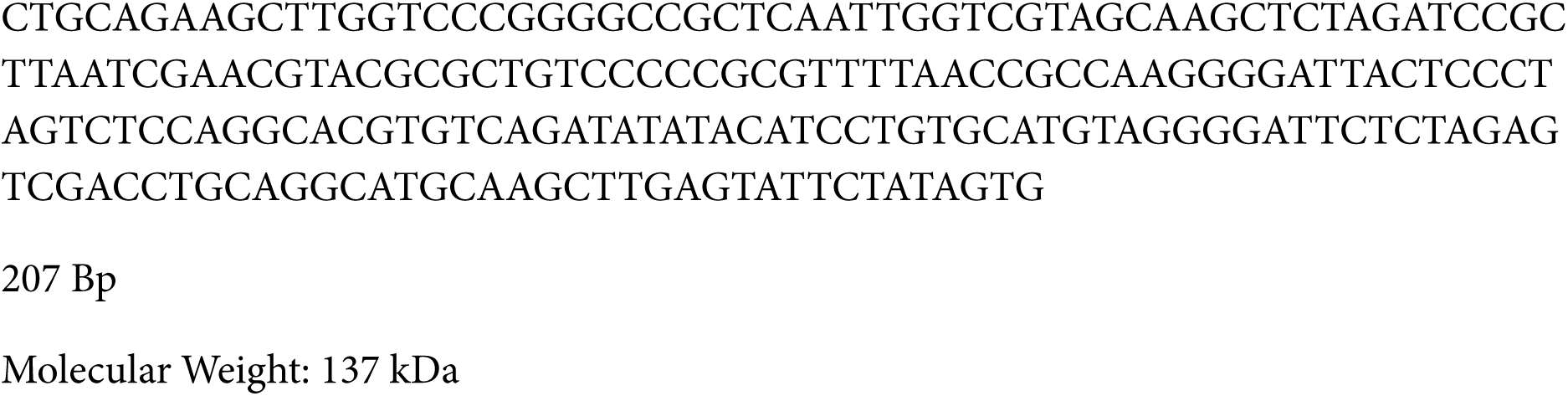
DNA sequence used in this study

