## Supplementary Videos for "DNA Partitioning Modulates Liquid-to-Solid Transitions and the Internal Microstructure of FUS Condensates"

### Slide 1
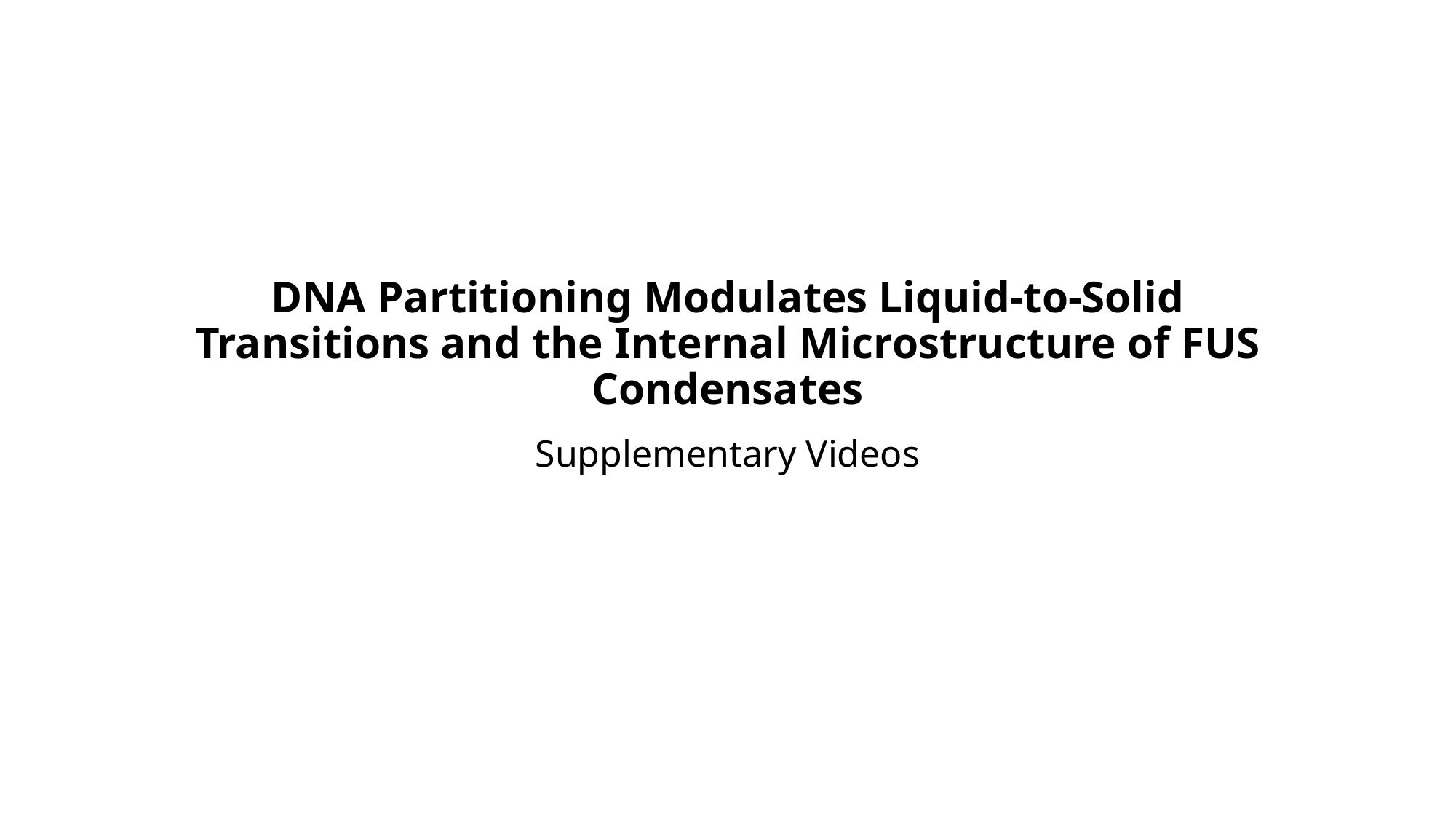

#### DNA Partitioning Modulates Liquid-to-Solid Transitions and the Internal Microstructure of FUS Condensates
Supplementary Videos

### Slide 2
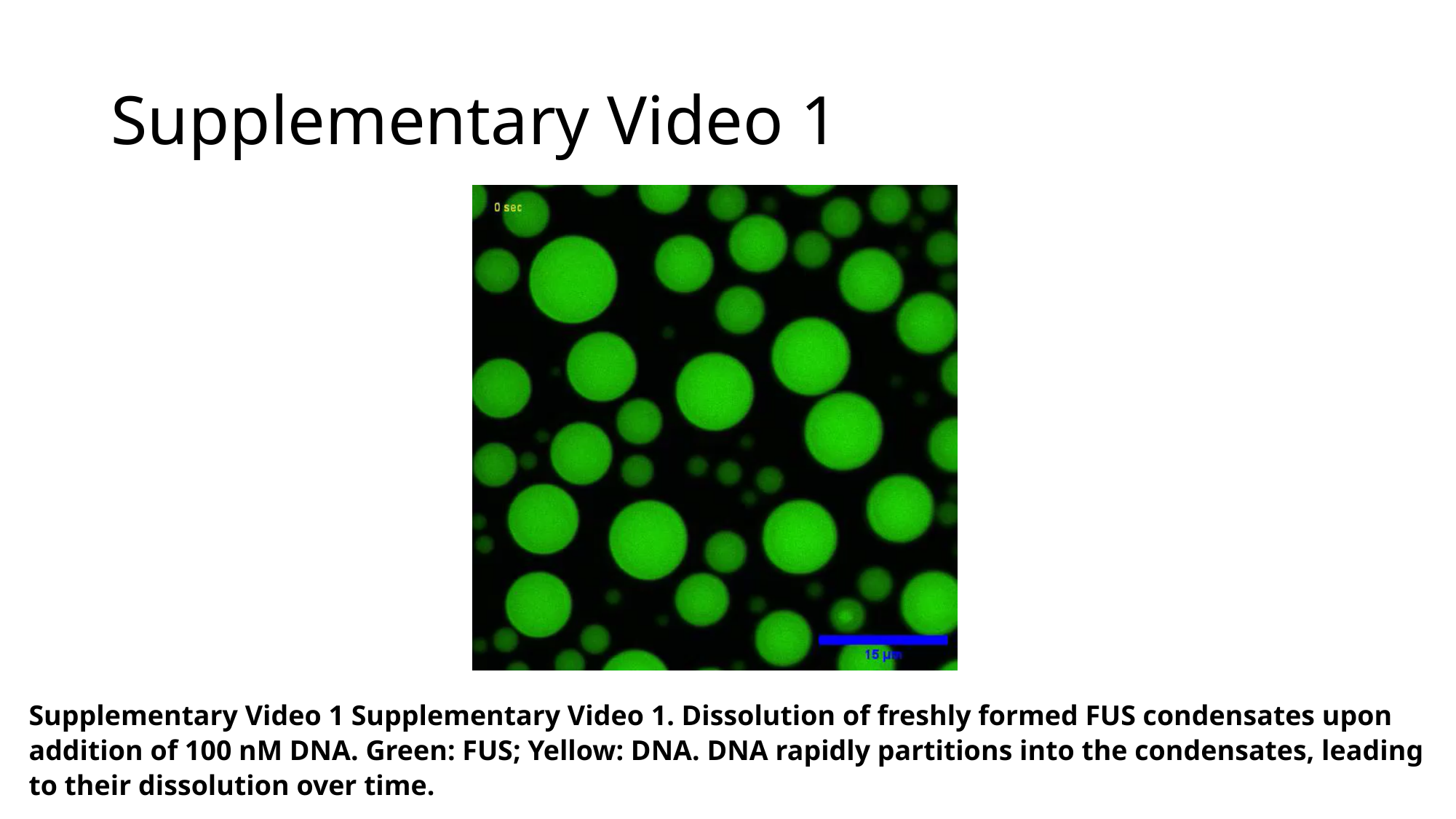

#### Supplementary Video 1
Supplementary Video 1 Supplementary Video 1. Dissolution of freshly formed FUS condensates upon addition of 100 nM DNA. Green: FUS; Yellow: DNA. DNA rapidly partitions into the condensates, leading to their dissolution over time.

### Slide 3
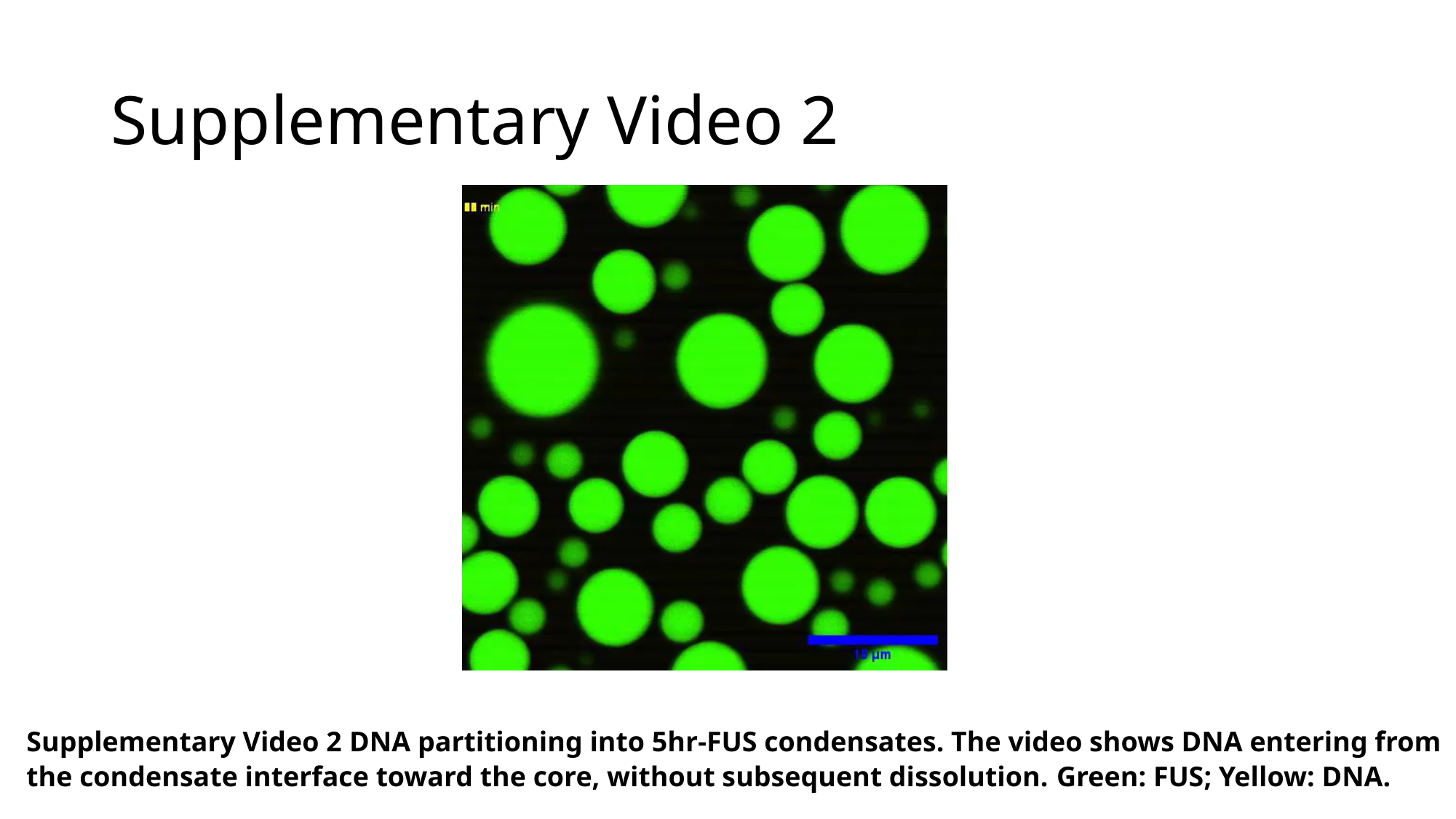

#### Supplementary Video 2
Supplementary Video 2 DNA partitioning into 5hr-FUS condensates. The video shows DNA entering from the condensate interface toward the core, without subsequent dissolution. Green: FUS; Yellow: DNA.

### Slide 4
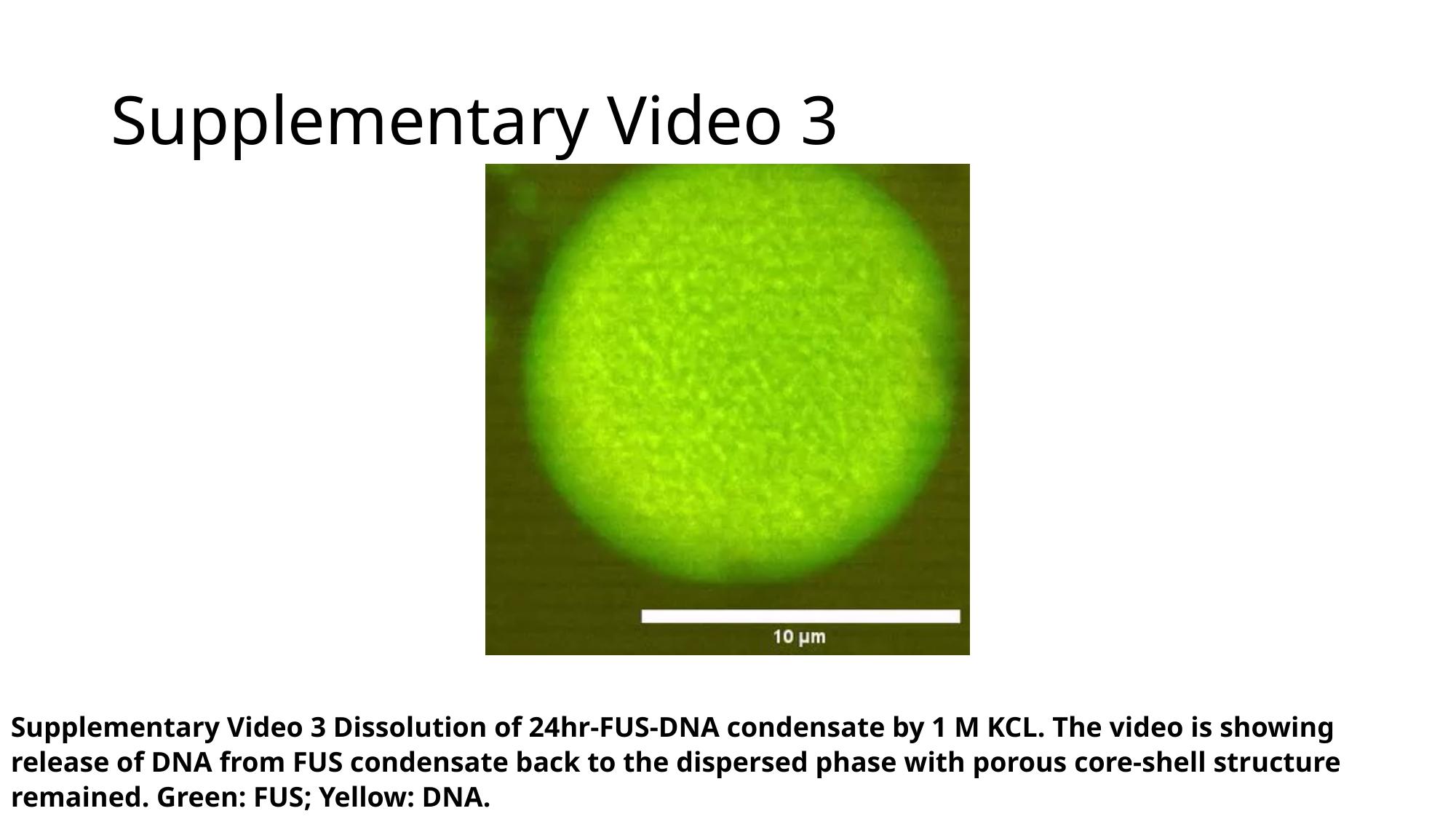

#### Supplementary Video 3
Supplementary Video 3 Dissolution of 24hr-FUS-DNA condensate by 1 M KCL. The video is showing release of DNA from FUS condensate back to the dispersed phase with porous core-shell structure remained. Green: FUS; Yellow: DNA.

### Slide 5
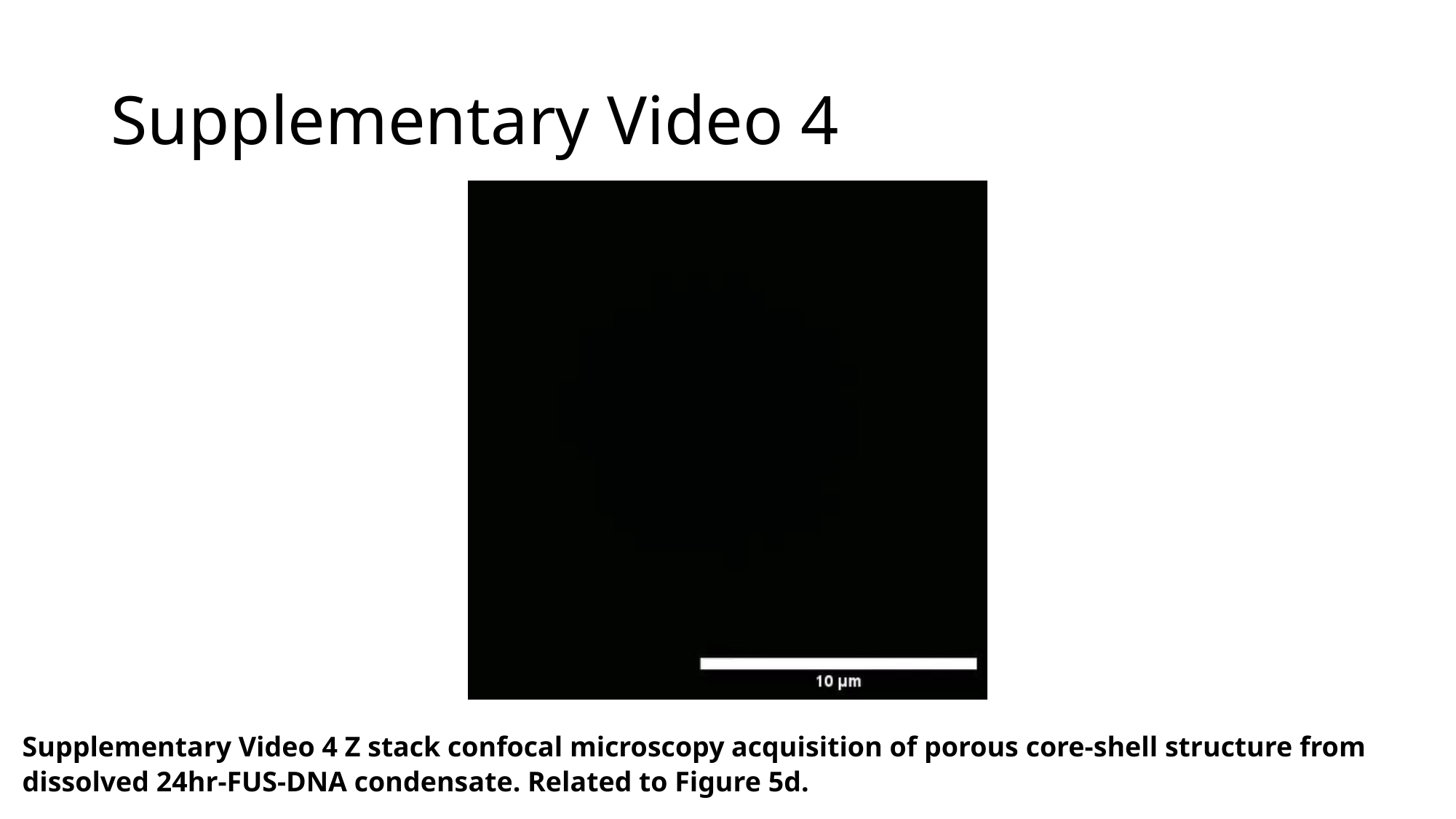

#### Supplementary Video 4
Supplementary Video 4 Z stack confocal microscopy acquisition of porous core-shell structure from dissolved 24hr-FUS-DNA condensate. Related to Figure 5d.

### Slide 6
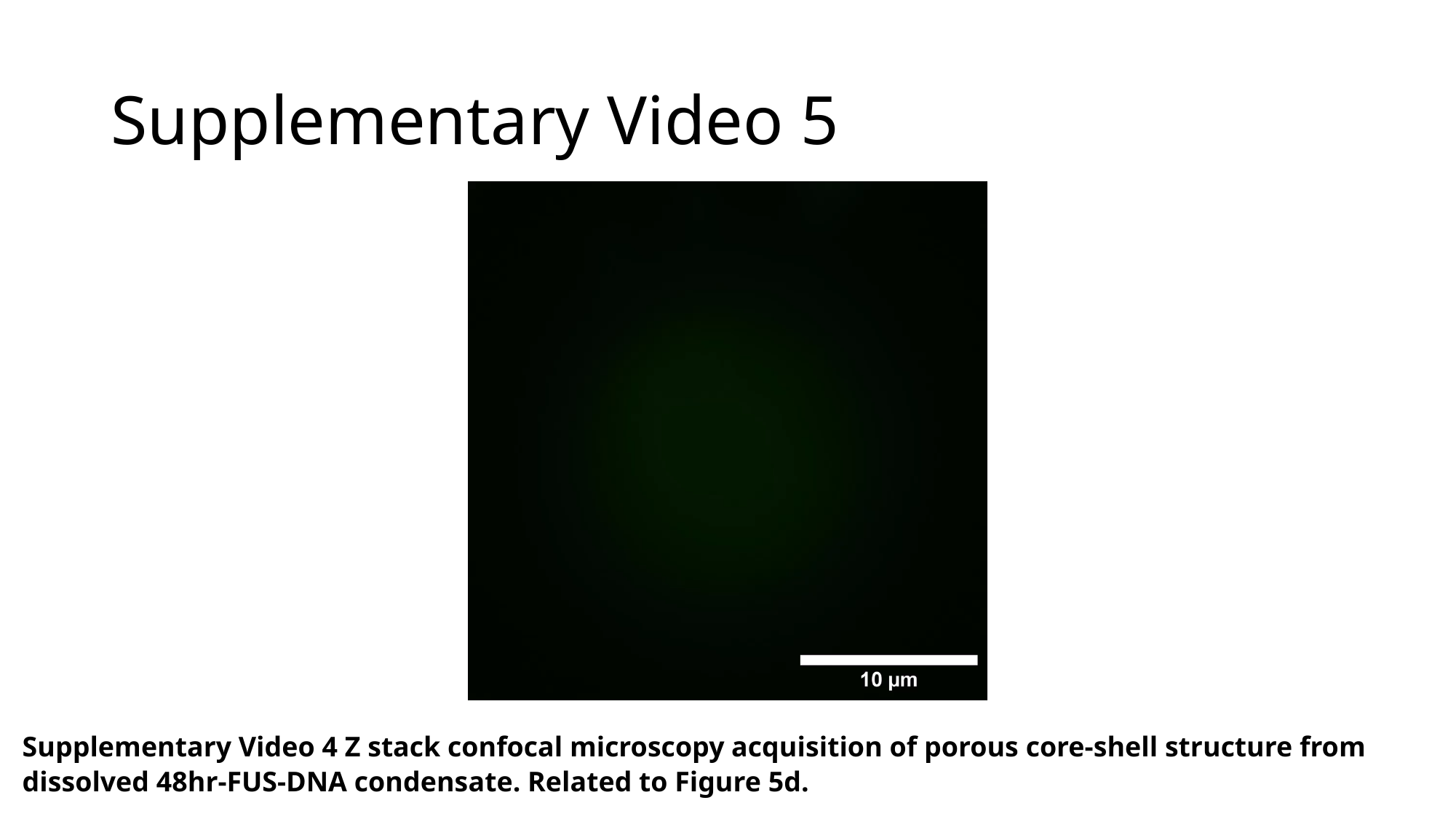

#### Supplementary Video 5
Supplementary Video 4 Z stack confocal microscopy acquisition of porous core-shell structure from dissolved 48hr-FUS-DNA condensate. Related to Figure 5d.

### Slide 7
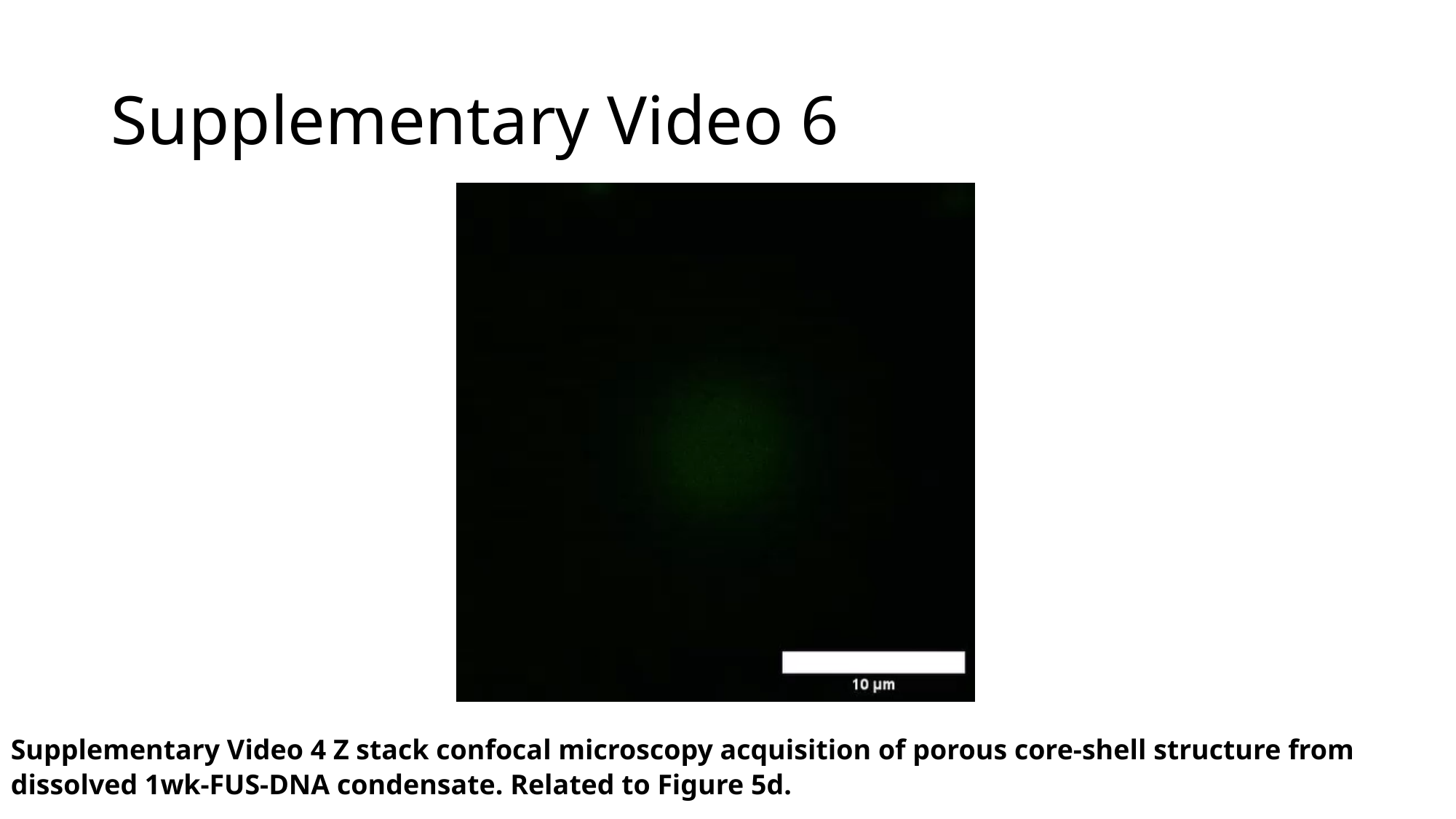

#### Supplementary Video 6
Supplementary Video 4 Z stack confocal microscopy acquisition of porous core-shell structure from dissolved 1wk-FUS-DNA condensate. Related to Figure 5d.
